# SLOGEN: A Structure-based Lead Optimization Model Unifying Fragment Generation and Screening

**DOI:** 10.1101/2025.10.14.682343

**Authors:** Bo Yang, Yunong Xu, Chijian Xiang, Yu Zhu, Tongtong Li, Anton V. Sinitskiy, Jianing Li

## Abstract

Lead optimization plays an important role in preclinical drug discovery. While deep learning has accelerated this process, structure-based approaches that leverage 3D protein-ligand information remain underexplored. Existing models could improve predicted affinity but often yield synthetically inaccessible compounds, whereas screening-based methods limit chemical novelty by relying on fixed fragment libraries. To bridge the gap, we introduce Slogen—a **S**tructure-based **L**ead **O**ptimization algorithm unifying fragment **G**eneration and scre**EN**ing. To achieve this, Slogen integrates a transformer-based variational autoencoder, pretrained on the BindingNet v2 dataset, with an E(3)-equivariant graph neural network that models 3D protein–fragment interactions. This unified framework enables both fragment generation and similarity-based screening, simultaneously addressing synthetic tractability and structural diversity. Benchmarking study shows that Slogen matches or surpasses state-of-the-art methods while exploring broader chemical space. Case studies on the Smoothened and D1 dopamine receptors demonstrate its capacity to design high-affinity, drug-like molecules, providing a practical method for structure-guided lead optimization.

## Introduction

Lead optimization—a crucial step in the preclinical phase of drug discovery—involves refining hit compounds to enhance their binding affinity, efficacy, selectivity, and pharmacokinetic/pharmacodynamic (PK/PD) properties. This process is notoriously expensive, labor-intensive, and time-consuming, yet essential before advancing candidates to clinical trials.^1, 2^ The classic lead optimization cycle includes designing and synthesizing compounds, testing properties of interest in relevant assays, and iteratively refining the structures based on feedback and data interpretation. Computer-aided drug design (CADD) methods—such as molecular docking^3^, alchemical free energy calculations,^4^ and quantitative structure–activity relationship (QSAR) modeling^5^—could help optimize hit molecules for higher affinity and improved drug-like properties, significantly accelerating the discovery process. In recent years, AI-aided drug design (AIDD) provides useful tools to nearly every stage of preclinical drug discovery.^6-8^ Advances in rational lead optimization with AIDD have been made possible with massive data and AI/ML technology, especially with deep learning.

Deep learning lead optimization algorithms can be broadly categorized into two types^9^: (1) goal-directed algorithms, which use reinforcement learning or other optimization techniques to design compounds and improve drug-like properties without structural constraints; and (2) structure-directed algorithms, which incorporate structural information to guide the generation of molecules or fragments using generative models. Since goal-directed lead optimization is inherently an optimization problem, traditional methods such as genetic algorithms, Bayesian optimization, and reinforcement learning are naturally applicable in this category.^9-11^ As a result, goal-directed approaches have been relatively well studied and applied across various lead optimization scenarios^12, 13^. However, these approaches are limited in their ability to leverage detailed three-dimensional structural information. Most rely on docking scores as a proxy for protein–ligand interactions, which provides only an indirect and often noisy representation of the true binding environment. In contrast, structure-directed strategies, which explicitly model protein–ligand structures, are more directly aligned with structure-based drug design (SBDD) but remain relatively underexplored.

Among various AI approaches, generative modeling has attracted significant attention for its ability to create novel molecules and scaffolds that could bypass known or patented compounds, thereby enabling the exploration of new chemical space.^9, 14^ Meanwhile, Recent advances in structural biology techniques— such as X-ray crystallography^15^ and Cryogenic electron microscopy (CryoEM)^16^—along with the emergence of structure-predictive models like Alphafold^17^ and RoseTTAFold^18^, have made structural information more accessible to both pharmaceutical companies and academic laboratories. This wealth of structural information, coupled with deep generative models, offers unprecedented opportunities for structure-based rational lead optimization.^19, 20^

However, compared to structure-based *de novo* generative algorithms that directly generate hit molecules within a binding pocket,^21-23^ structure-directed lead optimization algorithms— which aim to generate lead molecules with higher affinity than the starting compound using structural information—have received relatively less attention.^9^ While existing structure-directed lead optimization algorithms like Delete^24^ and DiffDec^25^ can generate molecules with potentially higher affinity, the synthetic accessibility of these newly generated compounds often limits their practical application. Although fragment-based algorithms such as DeepFrag^26^ partially address this issue by screening predefined fragment libraries, they inherently restrict the novelty and chemical diversity of generated candidates due to their dependence on fixed fragment sets.

To address the challenges outlined above, we developed **Slogen**—a **S**tructure-based **L**ead **O**ptimization algorithm unifying fragment **G**eneration and scre**EN**ing— which integrates fragment generation and virtual screening within a unified framework focused on fragment elaboration and growing tasks in lead optimization.^9^ (Figure 1) The framework integrates a transformer-based variational autoencoder (VAE), pretrained on fragments extracted from the BindingNet v2 dataset^27^, to learn a chemically meaningful latent space of fragment representations, enabling both generative decoding and similarity-based screening.. Additionally, an E(3)-equivariant graph neural network (GNN) is employed to capture the 3D structural context of the binding pocket and parent fragment to predict the latent vector corresponding to optimal elaborations.. From this predicted latent vector, diverse fragments capable of attaching to the parent scaffold and fitting within the binding pocket are sampled via the VAE decoder. The same latent vector can also be used to screen a predefined fragment library—encoded by the same VAE encoder—through cosine similarity–based fragment selection.

**Fig. 1.**
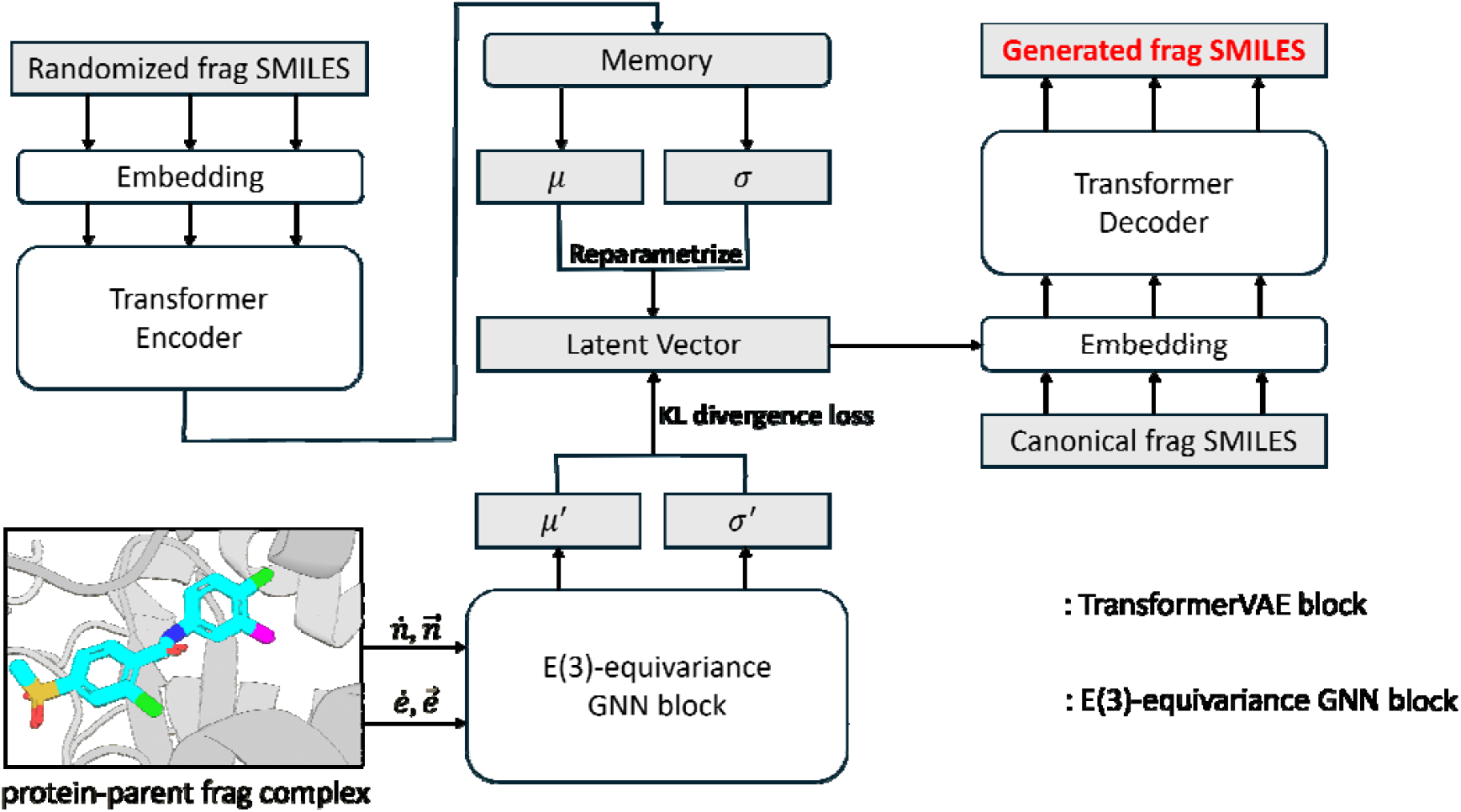
Overall structure of Slogen. The magenta atom indicates the attachment point of the given protein–parent fragment complexes. During generation, the latent vector is sampled from and predicted by GNN block. For screening, cosine similarity is calculated between the predicted and the vectors of a predefined fragment library encoded by the VAE encoder.

By training on a subset of BindingNet v2 dataset^27^, which offers potentially higher quality protein-ligand complexes than the commonly used CrossDock2020 dataset^28^, Slogen achieved comparable or superior performance in both generative and screening tasks across multiple metrics relative to state**-**of-the-art (SOTA) structure-directed lead optimization models such as Delete^24^ and DeepFrag^26^. Furthermore, the molecules generated by Slogen covered broader chemical space overall than fragments produced by other algorithms. Further case studies on the Smoothened receptor (SMO) and D1 dopamine receptor (D1DR) demonstrate the capability and flexibility of our model in real-world drug discovery applications. In summary, Slogen successfully addresses a major unmet need in structure-directed lead optimization—balancing synthetic feasibility with structural innovation—by coupling a learned chemical latent space with a 3D-aware predictive model. This unified framework bridges fragment generation and screening, providing a practical and scalable route toward structure-guided lead optimization.

## Materials and Methods

### Dataset

The BindingNet v2 dataset^27^, created by Zhu et al., contains 689,796 protein-ligand (PL) complexes extracted or modeled from PDB^29^ and ChEMBL^30^ databases. Due to computational resource constraints, we selected a high-confidence subset based on the hybrid score defined in the original study,^27^ resulting in 231,978 PL complexes. Each complex was further processed following a protocol similar to that used in DeepFrag,^26^ including the following steps:

1. Residues within 6 Å of the ligand were extracted to serve as the protein input.
2. Each ligand was split into multiple fragment pairs by iterating over all “cuttable bonds.” Dummy atoms were inserted at the cleavage points to mark the attachment sites on both fragments.
3. Only fragment pairs where the smaller fragment was less than 150 Da were retained.
4. The bond cleavage site had to be within 4 Å of the protein to ensure relevance to the binding pocket.

“Cuttable bonds” are defined as single bonds that are not part of a ring or aromatic system—i.e., bonds whose cleavage results in two separate fragments. For each ligand, all such bonds were iteratively cleaved to generate fragment pairs and saved as tuple in the format of (protein_6A.pdb, original_ligand.sdf parent_frag.sdf (big fragment), small_frag.sdf) for use in training stage. This exhaustive fragmentation strategy ensures comprehensive coverage of fragment space. While existing rule-based fragmentation methods such as RECAP^31^ and BRICS^32^ could account for synthetic accessibility of molecules, they typically yield less diverse fragments and fewer training pairs due to their reliance on predefined cleavage rules. Moreover, their limited flexibility can restrict applicability across diverse input complexes. In contrast, our iterative splitting approach generated a total of 1,795,968 fragment pairs, offering a rich and diverse dataset for training structure-directed generative models.

After fragment generation, we split the dataset into training and test sets using a UMAP-based clustering approach,^33^ applied to Protein–Ligand Extended Connectivity (PLEC) interaction fingerprints^34^ extracted from each PL complex. Unlike conventional sequence similarity-based splitting, this method accounts for cases where proteins may have low overall sequence similarity but share similar binding site environments—an important consideration for structure-based learning tasks. The final split resulted in 1,793,350 complexes in the training set and 2,618 complexes in the test set.

To evaluate the performance of our model in real-world drug discovery scenarios, we selected two hold-out test proteins with distinct characteristics. SMO (PDBid: 5l7i^35^) shares less than 30% sequence similarity with all proteins in the training and test sets. Additionally, its co-crystallized ligand was excluded from both sets. This setup represents an out-of-distribution test case, designed to assess the model’s ability to generalize to novel targets with minimal sequence homology. For another protein D1DR (PDB id: 7cky^36^), although D1DR itself is not included in the training or test sets, several proteins in the training set—such as the dopamine D3 receptor—share more than 40% sequence similarity. This scenario is intended to mimic real-world drug discovery settings, where researchers often work with targets that belong to known protein subfamilies and can leverage related structural or functional information. The inclusion of D1DR reflects this practical context.

### Overview of Slogen

Slogen is designed to perform fragment elaboration/growing given a 3D binding pocket and a parent fragment (Figure 1). We represent both the protein pocket and the parent fragment as a 3D k-nearest neighbor (KNN) graph, where nodes correspond to atoms and each atom is connected to its k nearest neighbors. Following Pocket2Mol,^21^ each node and edge includes both scalar and vector features, as incorporating both types enhances the representational power of the GNN.^21, 37^

Formally, the protein pocket 𝒫 is represented as a set of atoms 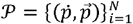, where 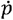 denotes scalar features of the *i*th heavy atom of the protein pocket and 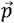 represents its vector feature of the th heavy atom of the protein pocket (i.e., 3D coordinates). Scalar features include atom element, amino acid type, backbone flag, and ligand atom flag (Table S1). Similarly, the parent fragment ℱ is also represented as a set of atoms 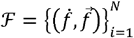, with 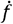 and 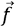 as the scalar and vector features of the th heavy atom in the parent fragment Scalar features for the parent fragment include atom element, ligand atom flag, attachment point flag, valence, and the number of different chemical bonds. Vector features are again the atomic coordinates. (Table S1). Following Pocket2Mol,^21^ we use a modified version of the Geometric Vector Perceptron (GVP) as an E(3)-equivariant GNN to aggregate information from the 3D protein–parent fragment complex. We denote the GNN here as *ϕ* and the input is processed as follows:

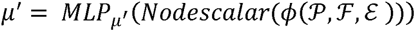

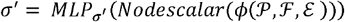

Here ℰ represents the edges in the KNN graph, which include the scalar features bond types and the distance encoded with Gaussian RBF kernels,^38^ and vector features unit directional vector of edges. The *Nodescalar* represents the E(3)-invariant scalar features of the attachment point atom. Fully connected layers are used to predict the vectors and *μ′, σ*′ which represent mean and standard deviation of a gaussian distribution and will be later used for both screening and generation.

The Variational Autoencoder (VAE) is a widely used generative model that has also been applied to molecular generation tasks.^39-41^ A VAE consists of an encoder–decoder pair that learns a latent representation of input data and can generate novel samples from a latent variable *z*. In addition to generation, the *μ* vector produced by the encoder can be used to screen a predefined library based on similarity.

To unify screening and generation within a single framework, we incorporate a VAE into our model. Specifically, we adopt the TransformerVAE developed by Yasuhiro Yoshikai et al.^42^ which combines the expressive power of transformers with the generative capabilities of VAEs. This model is particularly efficient for encoding and generating SMILES strings, which are sequential representations of molecules akin to natural language. The TransformerVAE is pretrained on small fragments (fragment below 150 molecular weight), which are extracted from the training dataset. The pretrained encoder can generate the *μ* and *σ* for each fragment, and diverse novel fragments can be generated by decoding the latent variable *z* sampled from the corresponding distributions. (Figure 1).

### Training process

A total of 38,987 small fragments, split from PL complexes, were converted into SMILES strings. The breaking points were marked using a dummy atom (*). These fragments were then used to pretrain the TransformerVAE model. Similar to a typical variational autoencoder (VAE),^43^ the model is trained by minimizing the negative Evidence Lower Bound (ELBO):^42, 43^

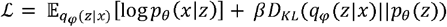

In a standard VAE, *β* is equal to one. However, to mitigate posterior collapse—often caused by the powerful transformer blocks—we set *β* to 0.01 during the training following TransformerVAE paper.^42^ Detailed model architecture and hyperparameters can be found in the original TransformerVAE paper.^42^

After pretraining, the TransformerVAE encoder can encode any small fragment into a latent space, producing vectors *μ* and *σ*. For each of the 1,795,968 protein–parent fragment complexes, we extracted the *μ* and *σ* vectors for ground truth fragments and used them as target outputs for training the GNN component. To improve training efficiency, only residues within 6 Å of the original ligand were extracted as the protein pocket input. Parent fragments were saved in .sdf format and used as input to the GNN with the protein pocket. The GNN was trained to minimize the KL divergence between its predicted Gaussian distribution (*μ*′ and *σ*′) and the ground truth distribution (*μ* and *σ*):

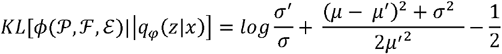

After 34 epochs, the validation loss converged. The checkpoint with the lowest validation loss was selected for downstream screening and generation tasks.

### Generation and Screening

To generate suitable fragments for a given protein–parent fragment complex, the complex is first processed through the GNN, which outputs two vectors: *μ*′ and *σ*′, representing the mean and standard deviation of a Gaussian distribution. A latent vector *z*′ is then sampled using the reparameterization trick:

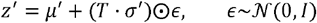

Here, T is a temperature parameter that controls the diversity of generated molecules. By default, T is set to one. The sampled latent vector *z*′ is then passed to the decoder of the pretrained TransformerVAE to generate a novel fragment.

For screening, the same GNN outputs *μ*′ and *σ*′. The predicted *μ*′ is used to screen a predefined fragment library, where each fragment has a corresponding *μ* vector extracted from the TransformerVAE encoder. Cosine similarity is computed between *μ′*and *μ* each in the library as follows:

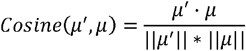

Fragments with higher cosine similarity to *μ*′ are considered more likely to improve binding affinity when attached to the parent fragments.

### Baselines

We compared our model with the following two state-of-the-art baselines:

- **Delete**.^24^ Delete is a GNN-based model trained using a masking strategy. It is capable of performing all four subtasks in lead optimization—namely, fragment elaboration, linker design, side-chain decoration, and scaffold hopping—in a 3D structure-based manner. In our study, we compared the performance of our model with Delete on the fragment elaboration (or growing) subtask, using the corresponding checkpoint provided in the GitHub repository.
- **DeepFrag**.^26^ DeepFrag is a 3D-CNN-based model that predicts the RDKFingerprint of the ground truth fragment given a 3D protein–parent fragment complex. The predicted RDKFingerprint is then used to screen a predefined fragment library based on cosine similarity. We compared our model with DeepFrag in terms of both screening and generation ability on the test sets for the fragment elaboration (or growing) subtask. However, since DeepFrag performs generation through screening, it cannot truly generate novel fragments.

### Evaluation Metrics

For the generation task, we generated 100 fragments for each protein–parent fragment complex in both our test set (Slogen test set) and the Delete test set using all three algorithms. For the screening task, we collected ground truth fragments for each test set to form two fragment libraries, and used Slogen, DeepFrag, and AutoDock Vina to screen the ground truth fragment for each protein–parent fragment complex in the two test sets. We evaluated generation and screening performance using the following metrics:

- **Vina score**. The Vina score estimates the binding affinity between ligands and the receptor. Since Slogen and DeepFrag output SMILES representations of fragments, we used the ‘scrub.py’ script from AutoDock Vina^44^ to generate ligand conformations by attaching the generated fragments to the parent fragments. We then performed docking using AutoDock Vina and recorded the Vina scores. More negative values indicate better predicted binding affinity.
- **QED**.^45^ The quantitative estimate of drug-likeness (QED) estimates the potential of a small molecule to become an oral drug based on the property distributions of approved drug molecules. The score ranges from 0 to 1, with higher values indicating better drug-likeness.
- **SA**.^46^ Synthetic accessibility (SA) score measures the ease of synthesizing a molecule, based on fragment frequency distributions from known compounds and structural complexity penalties. The score ranges from 1 to 10, with lower values indicating easier synthesis.
- **RO5**.^47^ Lipinski rule-of-5 (RO5) is a set of empirical rule that helps to predict if a biologically active molecule is likely to have the chemical and physical properties to be orally bioavailable. The rule includes: hydrogen bond donors <= 5. hydrogen bond acceptors <= 10. MW <= 500 and logP <= 5. The RO5 score ranges from 0 to 4, with higher values indicating that the molecule satisfies more of the rules.
- **Hit rate**. Hit rate represents the percentage of protein–parent fragment complexes for which the generated fragments exhibit higher predicted binding affinity (based on docking scores) than the original fragment when attached back.^48^ The value ranges from 0% to 100%, with higher values indicating the model’s ability to consistently generate fragments that improve binding affinity.
- **Better than original**. This metric indicates the percentage of generated fragments that, when attached to the parent fragment, result in better predicted binding affinity than the original ligand. The value ranges from 0% to 100%, with higher. values reflecting the model’s effectiveness in improving binding affinity through fragment generation.
- **Ring_freq > 100**.^20, 49^ This metric measures the percentage of ring systems in the generated molecules that occur more than 100 times in the ChEMBL ring system dataset. The value ranges from 0% to 100%, with higher values indicating better chemical plausibility and drug-likeness.
- **BM_ZINC20**.^20^ This metric evaluates the percentage of Bemis–Murcko (BM)^50^ scaffolds in the generated molecules that are also found in the BM scaffolds of drug-like molecules from the ZINC20 database. Higher values (0% to 100%) suggest better chemical plausibility and alignment with known drug-like structures.
- **BM_ZINC22**.^20^ This metric evaluates the percentage of Bemis–Murcko (BM)^50^ scaffolds in the generated molecules that are also found in the BM scaffolds of drug-like molecules from the ZINC22 database. Higher values (0% to 100%) suggest better chemical plausibility and alignment with known drug-like structures.
- **Top 1**,**5**,**10 Regeneration Rate**.^26^ Given a predefined fragment library containing the ground truth fragment for each protein–parent fragment pair, this metric measures the percentage of cases where the ground truth fragment is ranked within the top 1, 5, or 10 candidates selected by the algorithm. The value ranges from 0% to 100%, with higher values indicating better screening performance.

## Results

### Slogen achieves state-of-the-art generative performance in fragment elaboration/growing tasks

To evaluate its performance against other models, we randomly selected 261 protein–parent fragment complexes from the Slogen test set and generated 50 fragments per complex using each algorithm. For more comprehensive performance analysis, we also included the Delete test set, which contains 93 protein–parent fragment complexes. We then generated 100 fragments per complex for each algorithm. After attaching the generated fragments back to the parent fragments to create complete molecules (referred to as generated molecules), we computed various evaluation metrics.

Table 1 presents the generative benchmarking results on the Delete test set. Slogen achieves an average Vina docking score of –8.27 kcal/mol, outperforming both Delete (–8.07 kcal/mol) and DeepFrag (–7.52 kcal/mol). This score is also slightly better than that of the original ligands, indicating that Slogen can generate fragments that improve binding affinity when reattached. In terms of other affinity related metrics Hit rate and Better than original, Slogen achieves values of 92.1% and 43.8%, respectively, which are also the best performance among all tested models. For other metrics such as QED, SA, RO5 and Ring_freq > 100, Slogen ranks second among the tested models and performs comparably with the original ligands. This suggests that Slogen can generate molecules with favorable drug-like properties.

**Table 1.** Performance of the three algorithms across different metrics on the Delete test set. For each metric (excluding the original ligand), the top-performing result is highlighted in gray. Detailed explanations of each metric are provided in the Methods section.

|  | Original ligand | Delete | DeepFrag | Slogen |
| --- | --- | --- | --- | --- |
| Vina docking score (kcal/mol) (↓) | -8.09 | -8.07 | -7.52 | -8.27 |
| QED (↑) | $0.48 \pm 0.21$ | $0.46 \pm 0.21$ | $0.47 \pm 0.21$ | $0.46 \pm 0.20$ |
| SA (↑) | $3.58 \pm 1.21$ | $4.10 \pm 1.23$ | $3.66 \pm 1.12$ | $3.76 \pm 1.11$ |
| Ro5 (↑) | 3.41 | 3.37 | 3.55 | 3.39 |
| Hit rate (%) | - | 85.9% | 65.6% | 92.1% |
| Better than original (%) | - | 37.1% | 25.7% | 43.8% |
| Ring_freq > 100 (%) | 67.4% | 49.6% | 68.4% | 66.3% |
| BM_ZINC20 | 40.9% | 19.1% | 34.8% | 16.8% |
| BM_ZINC22 | 47.7% | 23.5% | 42.3% | 26.0% |

Figure 2A shows a t-SNE visualization of the chemical space coverage of fragments generated by different algorithms on Delete test set. Fragments generated by Slogen (points with blue color) and Delete (points with red color) occupy comparable overall areas but appear to span different regions of chemical space, likely reflecting differences in their fragmentation strategies. In contrast, fragments generated by DeepFrag (points with yellow color) cover a substantially smaller chemical space, consistent with the fixed library it employs.

**Fig. 2.**
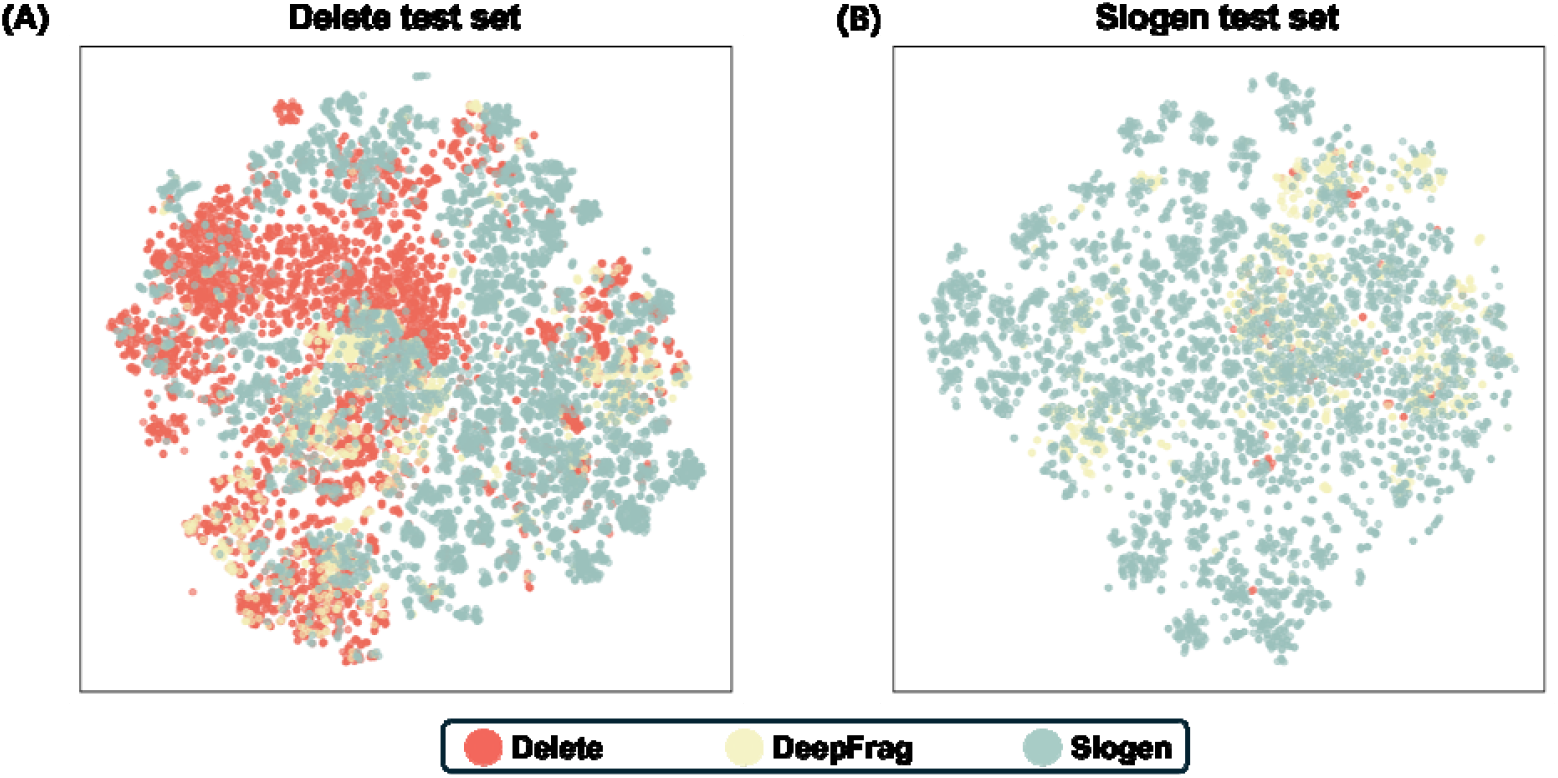
t-SNE visualization of fragments generated by each algorithm on the two test sets. Red, light yellow, and light green points represent molecules generated by Delete, DeepFrag, and Slogen, respectively. (A) Visualization for the Delete test set. (B) Visualization for the Slogen test set.

Table 2 summarizes the generative performance metrics on the Slogen test set. Consistent with the Delete test set results, Slogen achieved the highest Hit rate (91.8%) and “Better than original” percentage (55.0%), demonstrating its ability to improve the binding affinity of ligands of interest. It also exhibited improved synthetic accessibility, with the lowest average SA score (3.01), which was even lower than that of the original ligands (-7.62 kcal/mol). Although Slogen does not achieve the best Vina docking score on this set (-7.65 kcal/mol v.s. -7.81 kcal/mol from Delete and -7.52 kcal/mol from DeepFrag), its performance still surpasses that of the original ligands. The performance difference between the algorithms on Delete and Slogen test sets may be attributed to differences in fragment splitting methods. The Delete test set was generated using Delete’s own splitting strategy, which differs from the one used in our framework. Notably, for the Ring_freq > 100 metric, Slogen achieved a high occurrence rate of 80.1%, suggesting that it preferentially generates chemically reasonable ring structures frequently observed in ChEMBL and therefore potentially more drug-like and chemically plausible. Across other metrics, Slogen ranked second overall, with performance comparable to the leading algorithm.

**Table 2.** Performance of the three algorithms across different metrics on the Slogen test set. For each metric (excluding the original ligand), the top-performing result is highlighted in gray. Detailed explanations of each metric are provided in the Methods section.

|  | Original ligand | Delete | DeepFrag | Slogen |
| --- | --- | --- | --- | --- |
| Vina docking score (kcal/mol) (↓) | -7.62 | -7.81 | -7.52 | -7.65 |
| QED (↑) | $0.61 \pm 0.17$ | $0.60 \pm 0.18$ | $0.56 \pm 0.18$ | $0.57 \pm 0.19$ |
| SA (↑) | $3.06 \pm 1.00$ | $3.23 \pm 0.87$ | $3.24 \pm 0.99$ | $3.01 \pm 0.72$ |
| Ro5 (↑) | 3.90 | 3.80 | 3.83 | 3.76 |
| Hit rate (%) | - | 78.6% | 88.2% | 91.8% |
| Better than original (%) | - | 49.7% | 42.0% | 55.0% |
| Ring_freq > 100 (%) | 74.5% | 64.6% | 73.8% | 80.1% |
| BM_ZINC20 | 53.3% | 33.1% | 52.2% | 23.8% |
| BM_ZINC22 | 66.7% | 43.3% | 64.9% | 38.4% |

Figure 2 shows similar t-SNE visualization of fragments generated by different algorithms on the Slogen test set. Slogen covers substantially broader chemical space than DeepFrag and Delete. The weaker performance of Delete arises from differences in fragment-splitting strategies: binding pockets in the Slogen test set are generally smaller than those in the Delete test set, which restricts Delete to generate only a limited number of fragments for each protein–parent fragment complex.

In summary, Slogen demonstrated superior or comparable performance relative to SOTA models across multiple metrics on both test sets, with particularly strong results on affinity-related metrics, highlighting its ability to improve the binding affinity of ligands of interest. Furthermore, the t-SNE analysis shows that Slogen can be applied to diverse scenarios and covers broad chemical space without any deterioration in performance due to the fragment splitting strategy.

### Slogen demonstrates competitive performance with DeepFrag in fragment elaboration/growing screening tasks

Similarly, for generative tasks, we used both the Slogen test set and the Delete test set to assess screening performance. For each test set, the ground truth fragment corresponding to each protein– parent fragment complex was collected as a fragment library. Each model was then used to screen across the collected fragment library to determine whether it could retrieve the correct ground truth fragment. The screening results are summarized in **Table 3**.

**Table 3.** Comparison of screening ability between Slogen, DeepFrag and two base lines on Delete test set and Slogen test set. *r*.*r*. stands for regeneration rate.

|  | Vina docking | Slogen | DeepFrag | Random selection |
| --- | --- | --- | --- | --- |
| Delete test set performance |  |  |  |  |
| Top 1 r.r. | 4.7% | 20.4% | 7.5% | 1.1% |
| Top 5 r.r. | 9.4% | 30.1% | 15.1% | 5.4% |
| Top 10 r.r. | 16.5% | 35.5% | 20.4% | 10.8% |
| Slogen test set performance |  |  |  |  |
| Top 1 r.r. | 0.4% | 18.1% | 26.8% | 1.0% |
| Top 5 r.r. | 3.1% | 22.2% | 28.4% | 4.8% |
| Top 10 r.r. | 8.1% | 25.7% | 32.6% | 9.4% |

On the Delete test set, DeepFrag achieved Top-1, Top-5, and Top-10 regeneration rates of 7.5%, 15.1%, and 20.4%, respectively. In comparison, Slogen performed substantially better across all levels, achieving 20.4% for Top-1, 30.1% for Top-5, and 35.5% for Top-10 regeneration. Both models significantly outperformed commonly used methods such as AutoDock Vina and random selection,^26, 51, 52^ underscoring their ability to recover the ground-truth fragment given a 3D protein– parent fragment complex.

On the Slogen test set, however, Slogen underperformed relative to DeepFrag, achieving Top-1, Top-5, and Top-10 regeneration rates of 18.1% vs 26.8%, 22.2% vs 28.4%, and 25.7% vs 32.6%, respectively. Nonetheless, it still outperformed AutoDock Vina and random selection by a wide margin. This performance gap may be attributed to two key factors: 1. The Slogen test set was constructed using UMAP clustering followed by splitting, which creates a more challenging and diverse evaluation set.^33^ This makes it a more rigorous test of model generalization. And 2. DeepFrag is specifically designed for screening tasks, with a loss function tailored to optimize retrieval performance. This specialization contributes to its superior screening results, though it comes at the cost of weaker generative performance compared to Delete and Slogen (Tables 1 and 2, Figure 2). In summary, Slogen demonstrates screening performance comparable to DeepFrag and significantly superior to traditional methods such as Vina docking and random selection, underscoring its potential utility in real-world fragment-based drug discovery applications.

### Case study on SMO and D1DR shows the capability and versatility of Slogen on real-world drug design

We selected the Vismodegib–SMO complex (PDB ID: 5l7i) as the template for our case study, given that Vismodegib is an FDA-approved drug with multiple published SAR studies. As shown in Figure 3A, the terminal aromatic ring (highlighted in red) was removed from the Vismodegib structure, and Slogen was used to generate 100 fragments to replace this moiety, with the aim of improving both the Vina docking score and QED of the resulting molecules. For example, in the second structure of Figure 3A, Slogen replaced the aromatic ring with a tetrahydroisoquinoline ring, which improved both the docking score and QED value while maintaining a binding pose similar to that of Vismodegib. Beyond generation, as emphasized in the Introduction, screening capacity is also critical in real-world applications. We therefore screened against the DeepFrag fragment library to identify fragments capable of preserving or improving both the Vina score and QED value. As illustrated in the third structure of Figure 3A, replacing the aromatic ring with an indoline ring allowed Slogen to improve both the docking score and QED, thereby demonstrating its effective screening ability.

**Fig. 3.**
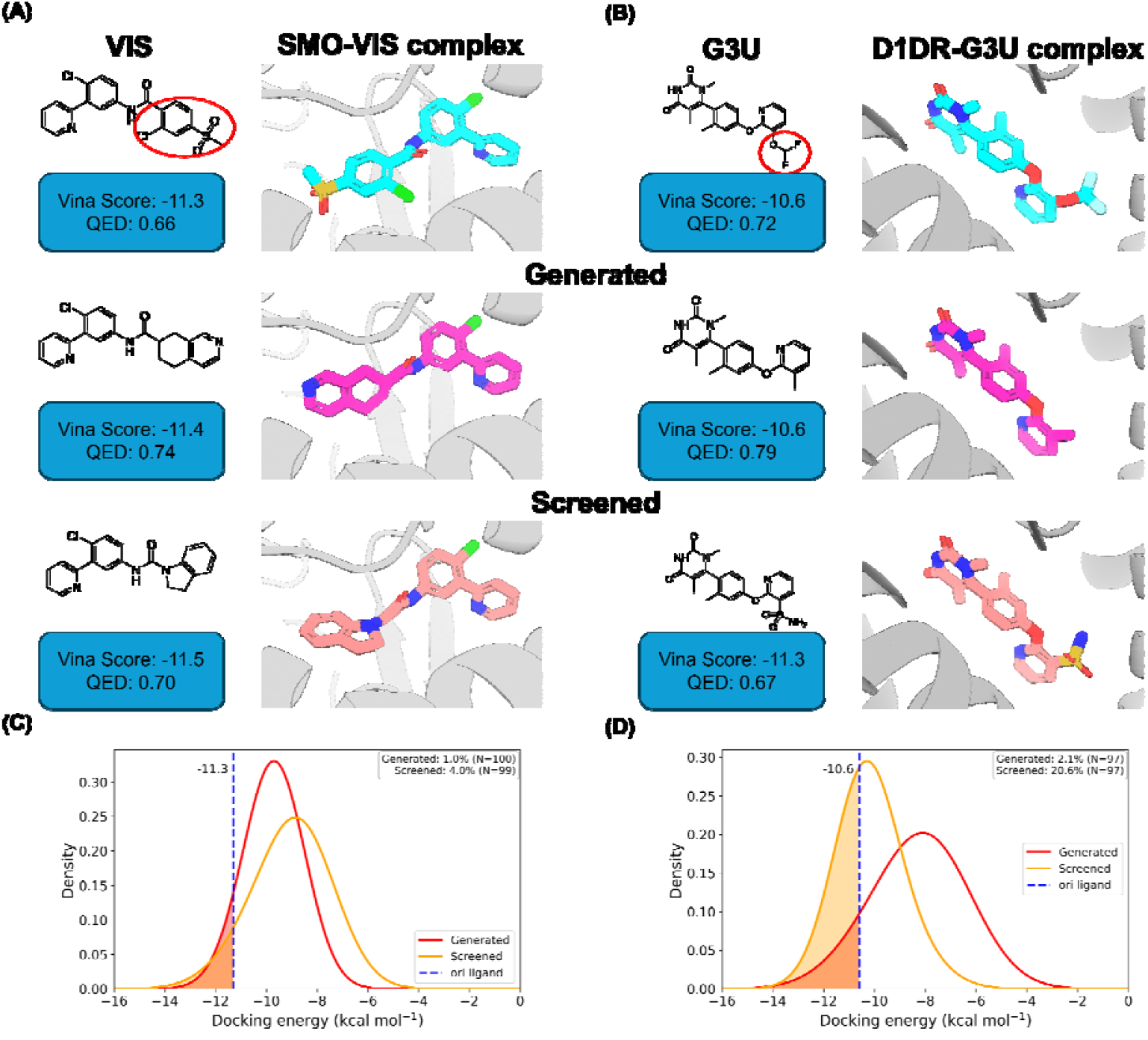
Case studies applying Slogen to Smoothened (SMO) and D1DR. (A, B) Slogen was applied to the SMO–VIS complex (PDB ID: 5l7i) and the D1DR–G3U complex (PDB ID: 7cky). The fragments circled in red were replaced using both the generation and screening functions of Slogen. The corresponding Vina scores and QED values of the generated molecules are shown. (C, D) Vina score distributions of molecules obtained from the generation and screening functions of Slogen. The Vina scores of the crystal ligands are indicated by blue dashed lines.

A similar trend was observed for D1DR. As shown in Figure 3B, Slogen successfully generated molecules with comparable Vina scores and higher QED values (second structures). When screening against the DeepFrag library, it produced molecules with substantially improved Vina scores while maintaining acceptable QED values (third structures). The Vina score distributions of generated molecules in these two case studies are shown in Figures 3C and 3D, where Slogen consistently produced molecules with better docking scores than the original ligands.

## Summary and Discussion

In this study, we developed Slogen, a 3D structure-based lead optimization model that unifies generation and screening to address the fragment elaboration/growing tasks in lead optimization. Benchmarking against Delete and DeepFrag on both the Slogen and Delete test sets demonstrates that Slogen is capable of generating diverse, high-affinity, and drug-like molecules from protein–parent fragment complexes, achieving SOTA performance in generative tasks. For screening tasks, although Slogen underperforms DeepFrag on the Slogen test set, it achieves comparable or even superior performance on the Delete test set. Moreover, Slogen significantly outperforms commonly used methods such as AutoDock Vina in terms of regeneration rates across both test sets, indicating its practical utility in real-world applications. Further case studies on SMO and D1DR show the capability of Slogen to generate molecules with better Vina score and QED value than reference molecules using both generation and screening functionality, which demonstrate the versatility of Slogen in unifying generation and screening.

Despite its strengths, Slogen has some limitations. In generative tasks, while it performs well on the Ring_freq > 100 metric, it performs poorly on the BM_ZINC20 and BM_ZINC22 metric. This suggests that Slogen effectively learns chemical priors related to ring structures and can generate chemically plausible rings but struggles to capture the distribution of BM scaffolds— particularly in the linker regions—when fragments are reattached to parent structures. To address this, several improvements can be considered: First, pretraining on a more diverse fragment set derived from drug-like molecules may help Slogen better learn chemical priors and generate higher-quality fragments. Second, incorporating a more powerful GNN architecture, as explored in recent studies,^53^ could enhance the model’s ability to extract meaningful features from protein–parent fragment complexes and guide more context-appropriate fragment generation.

In screening tasks, performance could be further improved by introducing a dedicated screening-specific loss function and then retraining the model, similar to DeepFrag, without altering the model architecture. This would allow generation and screening to use different checkpoints while remaining within a unified framework. Another idea could be applying contrastive pretraining to fine-tune the latent space, encouraging fragments with similar pharmacophores to cluster together while pushing dissimilar ones apart. This could enhance the model’s ability to retrieve relevant fragments during screening.

Additionally, we observed that fragmentation methods and selection of case studies significantly influence model performance. Current models like Delete, DeepFrag, and DiffDec use different fragmentation strategies, making direct comparisons difficult without retraining them on a unified dataset. For example, Delete performs better on its own test set—constructed using its native fragmentation method—than on the Slogen test set, which uses DeepFrag’s method. Besides, when applied to the D1DR task, Delete failed to generate a sufficient number of fragments due to the exclusion of small fragments (heavy atoms < 5) in its training data. Although we think retraining Delete using a more comprehensive fragment training set with corresponding binding pocket could solve this problem, this highlights the importance of a well-established, balanced fragmentation strategy that ensures both synthetic tractability and chemical diversity. For optimal real-world performance, input data should align with the fragmentation method used during model training, as the model is likely to have internalized those patterns. In future work, we aim to improve Slogen by incorporating the enhancements mentioned above, and develop a fragmentation method that better balances chemical intuition, diversity, and synthetic feasibility to support more robust and generalizable lead optimization. The code and data could be found at: https://github.com/Kartinaa/Slogen.

## Acknowledgement

We sincerely thank Dr. Jiaqi Guan and Haocheng Tang for their helpful discussions on this work. This work was supported by NIH R01 awards (GM129431/GM143370) and the AnalytiXIN fellowship (to J.L.).

