## Supplemental information for "SLOGEN: A Structure-based Lead Optimization Model Unifying Fragment Generation and Screening"

**Table S1.** Scalar and vector features included for protein atom, ligand atom and bonds during training Slougs.

| **Feature type** | | **Feature explanation** |
| --- | --- | --- |
| Protein atom node features | Scalar feature | Atomic element feature (C, N, O, S) |
|  |  | Amino acid type (20 nature amino acid) |
|  |  | backbone flag (0 or 1) |
|  |  | If ligand atom flag (all 0) |
|  | Vector feature | Atom coordinate |
| Ligand atom node features | Scalar feature | Atomic element feature (C, N, O, F, P, S, Cl, Br, I) |
|  |  | If ligand atom flag (all 1) |
|  |  | Total number of bonds (N_atoms, 1) |
|  |  | Sum of the bond types (N_atoms, 1) |
|  |  | If breaking point flag (0 or 1) |
|  |  | Number of bonds of each type (N_atoms, 3) |
|  | Vector feature | Atom coordinate |
| Edge feature for both ligand and protein | Scalar feature | Bond type (no_bond, single, double, triple) |
|  |  | Distance encoded with Gaussian RBF kernal |
|  | Vector feature | Unit directional vector |
